# BeEM: fast and faithful conversion of mmCIF format structure files to PDB format

**DOI:** 10.1101/2022.11.11.516190

**Authors:** Chengxin Zhang

## Abstract

Although mmCIF is the current official format for deposition of protein and nucleic acid structures to the Protein Data Bank (PDB) database, the legacy PDB format is still the primary supported format for many structural bioinformatics tools. Therefore, reliable software to convert mmCIF structure files to PDB files is needed. Unfortunately, existing conversion programs fail to correctly convert many mmCIF files, especially those with many atoms and/or long chain identifies. This study proposed BeEM, which converts any mmCIF format structure files to PDB format. BeEM conversion faithfully retains all atomic and chain information, including chain IDs with more than 2 characters, which are not supported by any existing mmCIF to PDB converters. The conversion speed of BeEM is at least ten times faster than existing converters such as MAXIT and Phenix. BeEM is available under the BSD licence at https://github.com/kad-ecoli/BeEM/.

## Introduction

The macromolecular Crystallographic Information File (mmCIF, also known as PDBx/mmCIF) format was introduced in year 2014 by the PDB database as its new standard for structure data deposition. The reason for the replacement of the previous official format (the legacy PDB format) by mmCIF is that all data fields in a PDB format file have fixed width, e.g., 5 characters and 1 character for an atom number and a chain identifier (chain ID), respectively. This limits the maximum number of atoms and chains in a PDB file to 99999 and 62, respectively. By contrast, the mmCIF format represents structure information as a space-separated tabular text file, where each data field can have unlimited length. This enables an mmCIF file to represent highly complicated structures with more atoms and chains than a PDB file. As of October 2022, for example, there are 3254 structures in the PDB database that are available as mmCIF but not as standard PDB format files.

Despite the advantages of mmCIF, for legacy reasons, the PDB format is still the only supported format for many bioinformatics applications ranging from side-chain packing (Huang, et al., 2020; Krivov, et al., 2009) and tertiary structure prediction (Zheng, et al., 2019) to structure alignment (Holm and Sander, 1995; Shindyalov and Bourne, 1998) and function prediction (Laskowski, et al., 2005; Zhang, et al., 2017). Even for some programs that support both mmCIF and PDB formats, PDB is still the preferred format due to smaller input size and faster file reading speed thanks to its fixed-width nature. For example, alignment of mmCIF structure by the TM-align program (Zhang and Skolnick, 2005) is twice as slow as aligning PDB structures.

To fulfill the need to use these programs on structures that are not available as a single PDB file, the PDB database provides “Best Effort/Minimal” PDB format, which splits a large mmCIF files into multiple smaller PDB files, each with up to 99999 atoms and up to 62 chains. A mapping file is also provided to map each original chain ID with two or more characters to a single-character chain ID in the split PDB file. The split PDB files and the mapping files are then bundled into a single TAR file. Despite its ability to encode arbitrarily large structures, there is not yet a publicly available webserver or standalone program for the generation of Best Effort/Minimal PDB files. Moreover, not all Best Effort/Minimal files provided by the PDB database are generated correctly, such as PDB ID: 7nwg, 7nwh, and 7nwi (Powers, et al., 2021).

To address this issue, several converters from mmCIF to PDB have been developed by the community (**Fig. 1A**). Among these conversion programs, BioPython (Cock, et al., 2009), cif-tools (https://github.com/PDB-REDO/cif-tools) and Atomium (Ireland and Martin, 2020) can only handle up to one character in chain ID, again limiting the number of distinct chains in the output PDB file to 62. MAXIT (https://sw-tools.rcsb.org/apps/MAXIT), GEMMI (Wojdyr, 2022) and Phenix (Liebschner, et al., 2019), on the other hand, handle two-character chain IDs in the output PDB files by occupying the usually unused column 21 in addition to column 22, the latter of which is reserved for the chain ID. They are, however, still unable to handle the 1036 structures from the PDB with chain IDs exceeding two characters.

**Figure 1.**
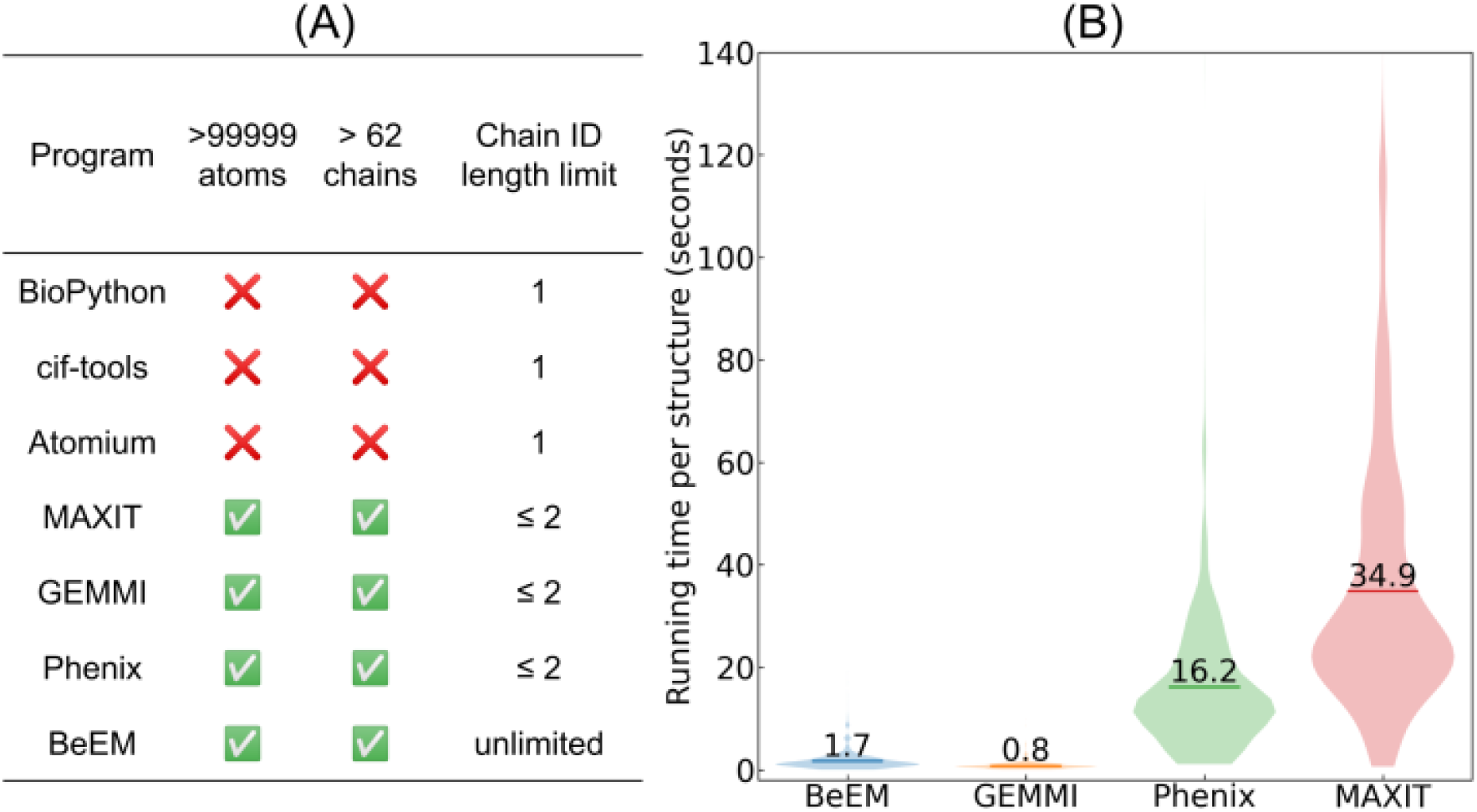
Comparison between BeEM and existing methods. **(A)** Limitations on the number of atoms and chains by different mmCIF to PDB conversion programs. Here, “Phenix” stands for the phenix.cif_as_pdb program from the Phenix package. **(B)** Running time of BeEM and three third-party programs for mmCIF to PDB format conversion. Horizontal bars indicate the average running time. BioPython, cif-tools and Atomium are not included because they cannot correctly generate PDB file for any of the input mmCIF structures.

To address these issues, this study proposed the <u>Be</u>st <u>E</u>ffort/<u>M</u>inimal (BeEM) program to convert mmCIF structure files to PDB files. It is currently the only open-source implementation for generation of Best Effort/Minimal PDB bundle files.

## Methods

BeEM is written in C++ without external dependencies, apart from the standard C++ 98 libraries. Following the Best Effort/Minimal file specification (https://www.rcsb.org/docs/general-help/structures-without-legacy-pdb-format-files), BeEM reads the _struct_keywords, _audit_author, _citation_author/_citation, _cell/_symmetry, _atom_sites, and _atom_site_anisotrop records from the input mmCIF files and outputs the HEADER, AUTHOR, JRNL, CRYST1, SCALE/ATOM/HETATM and ANISOU records in the PDB format files, respectively. Optionally, it can read the _entity_poly/_entity_poly_seq and _struct_ref/_struct_ref_seq records of the mmCIF files and convert them to SEQRES and DBREF records in PDB format, respectively. If the mmCIF input contains chain IDs with two or more characters, the user can choose to either output Best Effort/Minimal PDB files that map multi-character chain IDs to single character IDs or output a Phenix-style PDB file that retains two-character chain IDs.

## Results

BeEM, together with MAXIT, GEMMI, and Phenix, are benchmarked on a large dataset of 2218 structures from the PDB database that are available as mmCIF and Best Effort/Minimal files but not PDB format files. Although BeEM can handle any mmCIF format input, MAXIT, GEMMI and Phenix only handles up to two characters in the chain IDs. Therefore, only structures with up to two characters in their chain IDs are included in this dataset. On average, BeEM takes 1.7 seconds to convert an mmCIF file, which is slower than GEMMI but 9.6 and 20.7 times faster than Phenix and MAXI, respectively (**Fig. 1B**).

## Acknowledgements

The author thanks Dr Anna Pyle for insightful discussion and manuscript editing.

